# Characterization of Bilateral Reaching Development Using Augmented Reality Games

**DOI:** 10.1101/2024.06.05.597663

**Authors:** Shelby Ziccardi, Samantha Timanus, Ghazaleh Ashrafzadehkian, Stephen J. Guy, Rachel L. Hawe

## Abstract

**Highlights:**

- We developed augmented reality games to examine bilateral reaching in 133 children
- Symmetric and asymmetric reaching developed in parallel
- Synchrony in hands reaching targets improved with age
- Children continued to improve through age 17, though slower rate after 12 years
- Females demonstrated better reaching performance than males

Bilateral coordination is commonly impaired in neurodevelopmental conditions including cerebral palsy, developmental coordination disorder, and autism spectrum disorder. However, we lack objective clinical assessments that can quantify bilateral coordination in a clinically feasible manner and determine age-based norms to identify impairments. The objective of this study was to use augmented reality and computer vision to characterize bilateral reaching abilities in typically developing children. Typically developing children (n=133) ages 6-17 years completed symmetric and asymmetric bilateral reaching tasks in an augmented reality game environment. We analyzed the number of target pairs they could reach in 50 seconds as well as the time lag between their hands reaching the targets. We found that performance on both tasks developed in parallel, with development slowing but not plateauing after age 12. Children performed better on the symmetric task than asymmetric, both in targets reached and with shorter hand lags. Variability between children in hand lag decreased with age. We also found gender differences with females outperforming males, which were most pronounced in the 10-11 year olds. Overall, this study demonstrates parallel development through childhood and adolescence of symmetric and asymmetric reaching abilities. Furthermore, it demonstrates the ability to quantify bilateral coordination using computer vision and augmented reality, which can be applied to assess clinical populations.

## 1. Introduction

Bilateral coordination is essential for the majority of tasks of daily living (Bailey et al., 2015). One aspect of bilateral coordination is bilateral reaching, in which the arms perform symmetric or asymmetric reaching movements simultaneously, either to a single object such as a box, or separate objects such as a fork and knife. Such movements require the spatial and temporal coordination between limbs to successfully or most efficiently achieve the task goal, and therefore are more complex than unilateral reaching movements. Bilateral coordination is impacted in common disabilities in childhood including cerebral palsy (Decraene et al., 2021; Hawe et al., 2020; Smits-Engelsman et al., 2011; Utley & Steenbergen, 2006), developmental coordination disorder (Grohs et al., 2021; Huh et al., 1998), and autism spectrum disorder (Isenhower et al., 2012; Rodgers et al., 2018), which has significant functional ramifications due to the frequency of bilateral movements in everyday activities. In order to improve the assessment and treatment of bilateral coordination impairments, we first need to characterize the development of bilateral coordination in typically developing children. Additionally, we need quantitative assessment tools that can be deployed clinically without requiring specialized equipment.

The development of unilateral reaching in school-aged children and adolescents has been characterized using kinematic measures. Prior work has shown large changes in kinematic measures between 6 and 8 years old (Olivier et al., 2007). While some work has demonstrated that reaching kinematics achieve adult levels by ages 11 or 12 (Schneiberg et al., 2002; Simon-Martinez et al., 2018), others have showed performance does not plateau by 12 years, and continues to improve throughout teenage years (Gilliaux et al., 2015; Kuczynski et al., 2018; Olivier et al., 2007; Petuskey et al., 2007). While unilateral reaching abilities require only the coordination of one limb, interhemispheric processes are still at play as the contralateral limb’s activity must be suppressed.

Bilateral reaching abilities have been less characterized compared to unilateral reaches. Prior work has found that compared to unilateral reaching in 7-10 year olds, bilateral reaching showed less straight movements, indicating it may lag behind the development of unilateral reaching (Mazzarella et al., 2023). A comparison of unimanual and bimanual aiming movements showed that movement times for unimanual movements were faster than for bimanual movements up to age 8, and that symmetric movements were faster than asymmetric movements, however, only children up to 11 years old were investigated (Barral et al., 2006).

The developmental trajectory of bimanual coordination more generally is unclear and likely varies with different task demands. While not specifically probing reaching, two separate paradigms to examine bimanual coordination have found improvement in early childhood following a plateau at age 8, though (Riquelme et al., 2019; Serrien et al., 2014). An additional study found different timing for the development of temporal coupling, which consistently increased from age 6 through adulthood, while spatial coupling did not develop until adolescence (Boer et al., 2012). Heterogeneity in the paradigms used make it difficult to generalize developmental trajectories as tasks vary in whether they require symmetric or asymmetric movements between the limbs, as well as the conceptualization of the task goals and environmental influences. Additionally, prior work has been limited by small sample sizes and limited age ranges, which limit our ability to fully define the trajectory of bilateral reaching development.

Development of bilateral coordination is thought to correspond with the development of the corpus callosum (Jeeves et al., 1988; Marion et al., 2003; Rudisch et al., 2018). The maturation of the corpus callosum allows for the coordination of asymmetric movements between the limbs in addition to isolating unilateral movements. Early movements in babies and young children are often symmetrical, as isolating independent movement of each arm requires more mature coordination between the hemispheres of the brain (Brakke & Pacheco, 2019).

Mirror movements, in which voluntary movements in one limb are involuntarily mirrored on the contralateral limb, are present in typically developing children up to age 10, and thought to be abolished as the nervous system or more specifically the corpus callosum matures (Koerte et al., 2010). Studies that have examined interhemispheric transfer have found adult levels reached in older childhood to mid-adolescence, however structural differences are present between adolescence and adults signifying the corpus callosum matures into adulthood (Koerte et al., 2009). Interhemispheric transfer through the corpus callosum is thought to facilitate bilateral movements through the interhemispheric communication of feed-forward information, efference copies, and sensory feedback (Geffen et al., 1994). Additionally, sex differences in the development of the corpus callosum have been previously documented (Chavarria et al., 2014; Genc et al., 2018; Luders et al., 2010), which may translate into differences in development of bilateral coordination.

Assessment of reaching abilities have typically relied on 3-D motion capture or robotic devices in specialized laboratory environments. While kinematic characterization of reaching ability is needed, especially to determine spatiotemporal coordination between the limbs, the specialized equipment makes it difficult to translate assessments to clinical settings (Philp et al., 2022). Computer vision algorithms offer the opportunity to use readily available video, such as from a smartphone or webcam, to estimate pose information, or where the limbs are in space, and derive kinematic variables (Kidziński et al., 2020; Stenum et al., 2021). Computer vision has been used mostly for the analysis of gait, but has significant potential with upper limb movements as well (Cóias et al., 2022; Zamin et al., 2023). While the accuracy of computer vision approaches varies based on many factors including camera setup, plane of movement, and clothing, pose estimation algorithms have been shown to be valid, reliable, and accurate for many human motion analysis scenarios (Dill et. al., 2023; Latreche et. al., 2023).

The objective of this study was to characterize the development of bilateral reaching abilities in a large cohort of school-aged children and adolescents using computer vision and augmented reality games. Specifically, we examined performance on tasks in which children performed discrete reaching movements to pairs of targets requiring either symmetrical or asymmetrical reaching movements between the arms, probing coordination on tasks with independent goals (i.e. reaching separate objects) (Kantak et al., 2017). A secondary objective was to examine differences between bilateral reaching abilities in girls and boys, as previous work has found sex-based differences in the maturation of the corpus callosum (Chavarria et al., 2014; Genc et al., 2018; Luders et al., 2010). We hypothesized that reaching abilities would have the most rapid improvement prior to age 12 but continue to improve across adolescence. We also hypothesized that children would perform better on symmetrical reaches compared to asymmetrical reaches, and that symmetrical reaching ability would mature faster than asymmetrical. This study will improve our understanding of the development of bilateral coordination from ages 6 to 17 years through the uses of a large cohort, quantitative measures of symmetric and asymmetric reaching, and exploratory analyses of gender differences.

Additionally, the findings of this study will demonstrate a novel method of quantifying bilateral reaching abilities and provide normative data to assess impairments in children with neurodevelopmental conditions.

## 2. Methods

### 2.1 Participants

We recruited and enrolled typically developing children during the 2023 Minnesota State Fair at the University of Minnesota’s Driven to Discover Research Facility across 2 days. This setting enabled us to rapidly recruit from an approximately representative population and to sample from populations that might not otherwise participate in university-based research. Our inclusion criteria were: age 6-17 years, normal or corrected to normal vision, written informed consent (ages 16-17; or parental permission and child assent for ages 6-15). Our exclusion criteria were: any disorder or injury that limits the range of motion or otherwise changes the way the participant moves or controls their arms (based on participant/parent report), inability to comply with the study protocol. Participants and/or their guardians completed demographic questionnaires including age (in years), gender, handedness (defined as hand they write with), and questions on their video game experience (frequency of playing and familiarity with motion capture-based games). Participants also provided feedback on their experience with the study game after completion. Surveys and instructions were offered in either verbal or written English. The data collection session for each participant took approximately 10 minutes. This research was approved by the Institutional Review Board (IRB) at the University of Minnesota. All participants received a University of Minnesota branded backpack, valued at approximately $2, as an incentive.

### 2.2 Experimental Procedure

Participants were seated in an arm-less, stationary chair in front of a green screen facing a 32 in, 2560 x 1440 resolution computer monitor with a 720p Logitech C720 webcam mounted at the top of the screen. All graphics related to the tasks are shown on the computer monitor in front of the participant. The distance between the monitor/webcam and the participant’s chair was adjusted so that the participant could reach the entire tracked field displayed on the monitor with their shoulders abducted to 90 degrees (side reach) or flexed to 180 degrees (overhead reach). This scaled all reaches to approximately the participant’s arm length.

Bilateral reaching abilities were assessed using a custom-developed, gamified method of collecting kinematic information during reaching tasks. The game software, utilizing a single camera, tracks participants’ movements (sampled at approximately 60 frames/second) and allows them to interact with virtual targets displayed on a monitor in front of them. The game software was developed using the Unity 3D game development engine (version 2021.3.16f1) with MediaPipe (Lugaresi et al., 2019) for human pose estimation. Participants’ interaction with game objects is determined by their current pose as estimated from MediaPipe. Only the position of the hand in the frontal plane is used to interact with the game and the basis for the results in this current study.

Participants completed three different target reaching tasks within our gamified video-based assessment modality: a symmetric bilateral reaching task, an asymmetric bilateral reaching task, and a unilateral reaching task in which participants could select which hand to use for each reach. This manuscript focuses on the symmetric and asymmetric bilateral reaching tasks; results from the unilateral reaching task are not included in this analysis. Tasks were completed in a randomized order for each participant. Screenshots of game play for the symmetric and asymmetric tasks are shown in Figure 1. In each of these tasks, participants viewed themselves reflected on the computer monitor in front of them. This view was overlaid with two virtual reaching targets shown as stylized anthropomorphic squares, each with size approximately 5% of the play space. Two virtual targets were positioned to be either mirrored across the midline requiring symmetrical arm movements (Figure 1A, 1C) or asymmetrically positioned to require asymmetrical arm movements (Figure 1B, 1D). Participants were asked to “collect” the targets by moving their arms so that the hands of their digital reflection reached the displayed targets. When the MediaPipe hand tracking algorithm detects the average position of the four tracked points within each of the participant’s hands has overlapped their associated targets’ bounds for five frames consecutively (approximately 0.1 seconds), the target pair is considered collected and is replaced by a new pair of targets in a different location. Using the average position of multiple track points in the hand and the consecutive frame requirement decrease the influence of any tracking errors such as single frame tracked position jumps. In the development of the game, we extensively tested the system with a wide range of participants and conditions and confirmed the accuracy of detecting hand overlaps with the displayed targets based on visual observations of gameplay.

**Figure 1.**
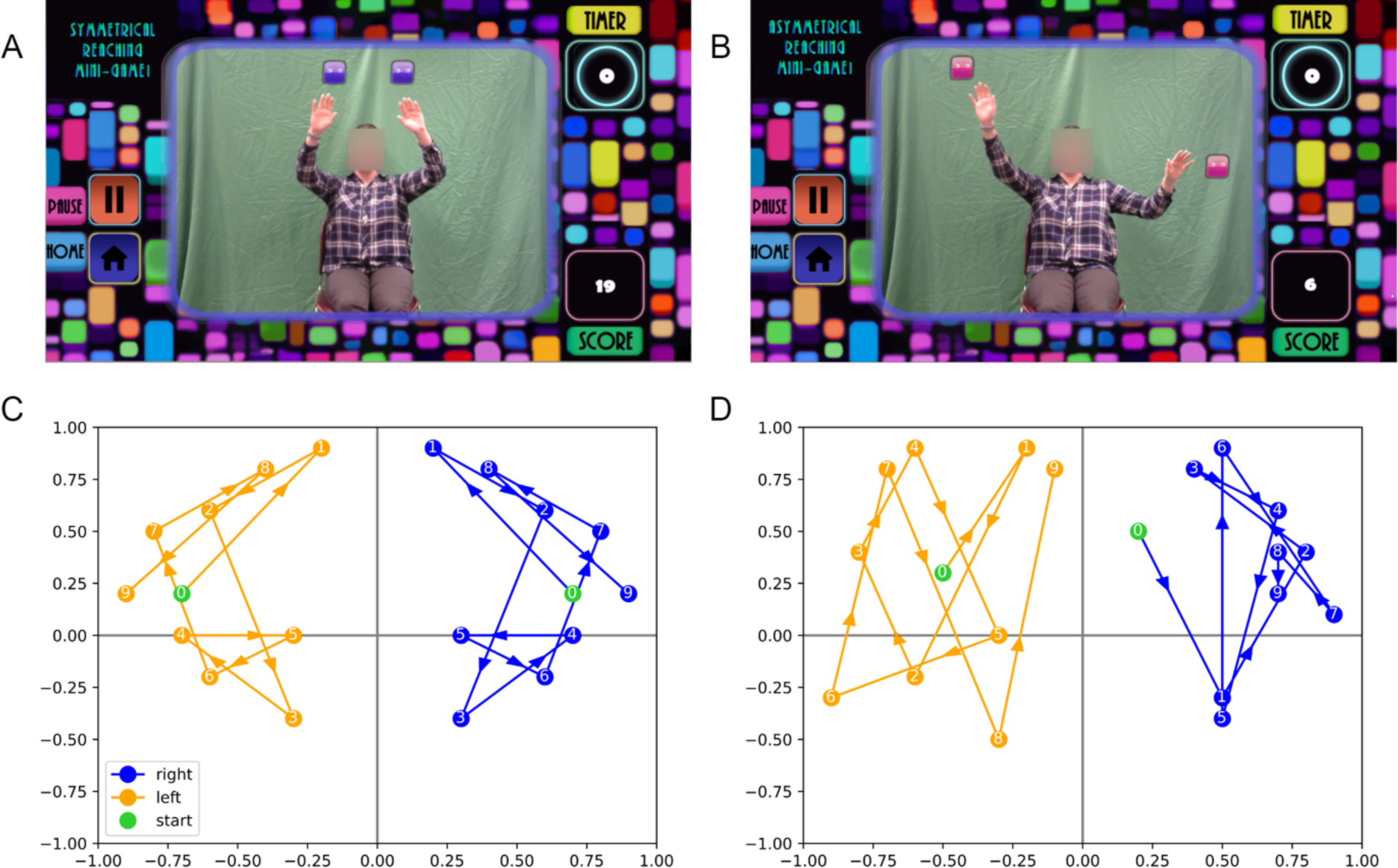
Screenshots of the developed game software. Screenshots are shown during gameplay for the symmetrical reaching task **(A)** and the asymmetrical reaching task **(B)**. Participants see themselves as well as two virtual targets (anthropomorphized squares) in the center. The screen also displays the countdown timer and score on the right side of the screen, with game control buttons on the left side of the screen. Additionally, a map of the positions and direction of movement between each pair of targets with pre-determined locations and order (the first ten targets) for the symmetrical reaching task **(C)** and the asymmetrical reaching task **(D)** are shown. The axes represent the center of the game space with the origin approximately matched to the center of the participant’s chest in calibration.

In each task, the participants completed an untimed practice round where they collected five pairs of practice targets and could ask questions. Participants then completed a gameplay round where they were asked to collect as many targets as possible within 50 seconds (a numeric and a visual timer were displayed on screen). This time frame was selected after pilot testing to minimize fatigue and maintain attention while ensuring we could assess enough reaches within the time period to sample the workspace. Participants received one point for each target pair they collected, with the total points (or score) displayed in the corner of the screen. The first ten pairs of targets in each task had predetermined locations that were chosen to span the workspace (positions illustrated in Figure 1C and 1D). The next ten pairs of targets had positions randomly selected from the list of ten predetermined locations. Any targets the participant collected past 20 in number were randomly placed around the screen based on the task parameters (symmetrically or asymmetrically located) and at least 300 pixels distance away from the previous target location. We aimed to have targets distributed across the workspace and comprising a range of movements in terms of distance and direction, as well as varying degrees of asymmetry both in the distance and direction. This would allow us to collect information that can generalize across different movements. Additionally, we also selected positions so that it would be intuitive which hand to use for which target based on the positioning. Across all participants, the average distance from one target to the next within each task was not statistically different between the symmetric and asymmetric bilateral reaching tasks (*t(134)=0.031, p= 0.975*).

### 2.3 Primary Outcomes

We use the total number of pairs of targets (i.e., score) collected in each 50 second reaching task as the overall measure of task performance on the symmetric and asymmetric reaching tasks. We also use the absolute value of the time difference between when each hand reached its associated target as a measure of synchronization of movement, which we refer to as hand lag. The unsigned value (absolute value) was used as target location and distance may influence which target was reached first. For each participant, we used the median hand lag across all reaches to be robust against reaches with uncharacteristic values. We also examined for each individual what percentage of target pairs they reached first with the dominant hand.

### 2.4 Statistical Analysis

The data from any participant who did not complete both the symmetric and asymmetric reaching tasks was removed (4 removed; n = 133). In rare cases, participants would continue moving their hands after target collection and the next target would generate overlapped with their new hand position and thus were unintentionally collected. In order to only consider intentionally collected targets, all targets collected less than 0.25 seconds after their appearance were assumed to be unintentionally collected and were removed from the total score and hand lag measures. All participants whose score or median hand lag exceeded four quartiles away from the associated mean were considered outliers and removed from the associated analysis (3 removed from hand lag analysis).

Normality was assessed independently on scores from the symmetric and asymmetric bilateral reaching tasks with a Kolmogorov-Smirnov Test and it was found that both can be considered normally distributed (symmetric reaching task *D(133)* = 0.073, *p* = 0.449; asymmetric reaching task *D(133)* = 0.09, *p* = 0.218). Parametric analysis was therefore used for the performance measure. The normality of median hand lag was similarly assessed and it was found that median hand lags of the symmetric task can be considered normally distributed (*D(130)* = 0.146, *p* = 0.176), however the median hand lag measures from the asymmetric reaching task may not be normally distributed (*D(130)* = 0.193, *p* = 0.036). Thus, non-parametric analysis was used for the hand lag measures.

To examine how task performance develops with age, we first fit a second-order polynomial curve to the participants’ total score as well as investigate the development of synchrony by fitting a similar curve to the median hand lag in each task across the age range. To test our hypothesis of continued development after 12 years of age, we perform a split-half analysis by fitting separate lines to scores of participants less than 12 years old and 12 years and older. By analyzing the slope of the fit for children over 12 years, we determine if development continues and how the relative speed of their development compares to that of earlier ages.

To examine the overall difference in performance and temporal synchrony between the symmetric reaching task and the asymmetric reaching task, we perform a paired t-test of scores and a Wilcoxon Signed-Rank Test on median hand lag values. To determine if the differences in performance and temporal synchrony of symmetrical and asymmetrical reaching change (widen or narrow) across development, we also conducted regression analysis on the difference between the two tasks’ scores and median hand lag across age.

Additionally, we examined whether there were gender differences in performance by first using an independent t-test. Since gender effects may also be different across the age span, we also used a two-way ANOVA to examine the effect of both age and gender on task performance, first binning participants by ages with approximately equal numbers of participants. The age categories were as follows: 6-7 year olds, 8-9 year olds, 10-11 year olds, 12-13 year olds, and 14-17 year olds. The larger age span for 14-17 year olds accounted for lower numbers of teens enrolled and also reflects the less expected development during adolescence. Additionally, post-hoc independent t-tests were performed between the scores of male and female participants within each age group.

Lastly, we explored the effect of possible confounding factors on task performance, namely video game experience and motion capture experience, as obtained by our questionnaire, by performing an ANCOVA test on symmetric reaching task performance and asymmetric reaching task performance between different video game and motion capture experience levels while controlling for participant’s age.

We used Python 3 with the scipy, statsmodels, and sklearn libraries for all statistical analysis.

## 3. Results

### 3.1 Demographics

We recruited and enrolled 151 total participants, of which 133 participants had complete data from both the symmetric and asymmetric tasks and were included in data analysis. The age and gender distribution can be seen in Figure 2. Individuals who reported non-binary / third-gender or did not report a gender were not included in gender-based analyses due to small sample size (n=4). The binned age groups used for gender analysis discussed in Section 2 have populations of approximately equal size: 6-7 year olds (*n = 19*), 8-9 year olds (*n = 26*), 10-11 year olds (*n = 34*), 12-13 year olds (*n = 25*), and 14-17 year olds (*n = 25*).

**Figure. 2.**
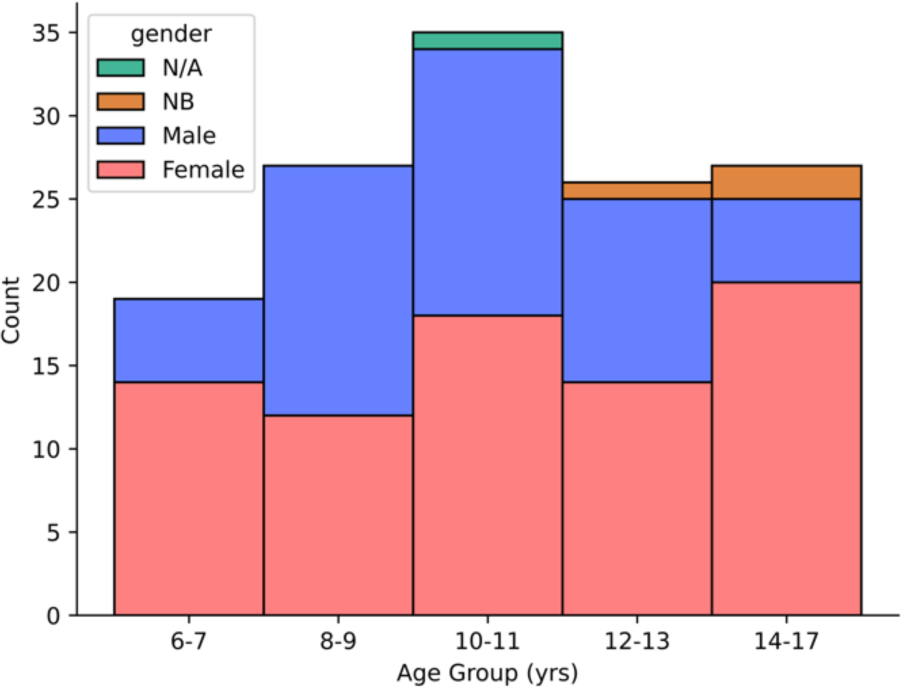
Total age and gender distribution. Distribution of participants, segmented by reported gender (female (red), male (blue), non-binary/third gender (orange), and unreported (green)), included in each defined age group.

### 3.2 Performance Development with Age

Figure 3A. shows scores on each task across the age range, as well as the quadratic curves fit to the data. The quadratic fits demonstrated that both symmetric bilateral task performance (*R^2^* = 0.512, *p* = 0.004), and asymmetric bilateral task performance (*R^2^* = 0.518, *p* < 0.001) improved with age, with age explaining over half of the variance in performance.

**Figure. 3.**
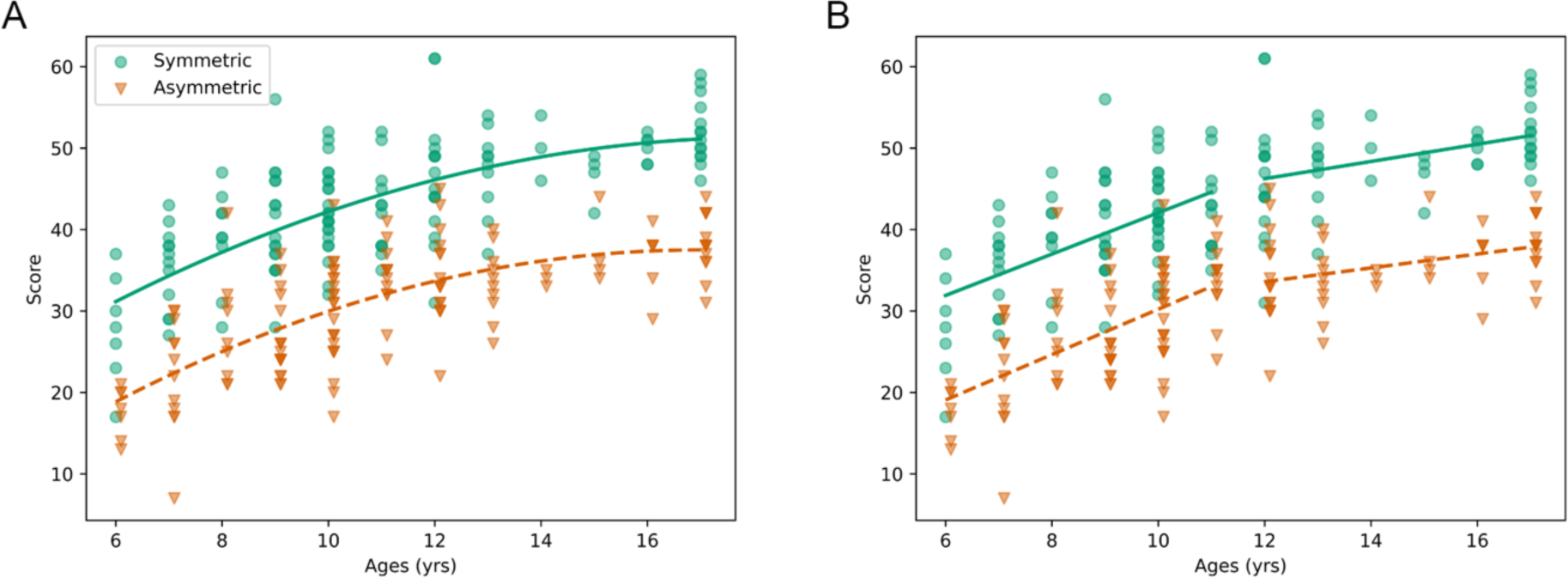
Development curves for total score. **(A)** Participants’ score in the symmetric reaching task (green circle) and the asymmetric reaching task (orange triangle) are plotted against the participants’ age. Quadratic curves are fit to the data for each task (symmetric task, solid line; asymmetric task, dashed line). **(B)** Split regression analysis is performed on the same data, with separate linear fits for 6-11 year olds and 12-17 year olds. Darker colors indicate overlapping data points.

Split-half regression analysis was done to compare the relative rates of change before and after age 12, as shown in Figure 3B. We find that performance continues to increase after the age of 12 (symmetric reaching task: *p* = 0.005, *slope* = 1.058, 95% C.I. [0.339, 1.778]; asymmetric reaching task: *p* = 0.006, *slope* = 0.853, 95% C.I. [0.254, 1.451]) but at a slower rate than before the age of 12 (symmetric reaching task: *p* < 0.001, *slope* = 2.538, 95% C.I. [1.686, 3.389]; asymmetric reaching task: *p* < 0.001, *slope* = 2.785, 95% C.I. [1.965, 3.605]).

We found median hand lag decreased with age for both the symmetric (*R^2^* = 0.312, *p* < 0.001), and asymmetric (*R^2^*= 0.283, *p* < 0.001) reaching tasks. Hand lag development curves for each task can be seen in Figure 4. Figure 4 also demonstrates how variability between children also decreases with age in both tasks, with the standard deviation of median hand lag values of the 6-7 year olds being 42 ms in the symmetrical task and 182 ms in the asymmetrical reaching task and both decreasing to a standard deviation of 12.5 ms and 51.2 ms respectively for the 14-17 year olds. This equates to a 70.3% decrease in variability in the symmetric task and 71.9% decrease in the asymmetric task across our collected age span. As the differences in the scale of hand lag between the two tasks makes it difficult to compare the developmental trajectories between tasks visually, as seen in Figure 4A, we also plotted the development curves using the normalized (z-score) of hand lag, as shown in Figure 4B.

**Fig. 4.**
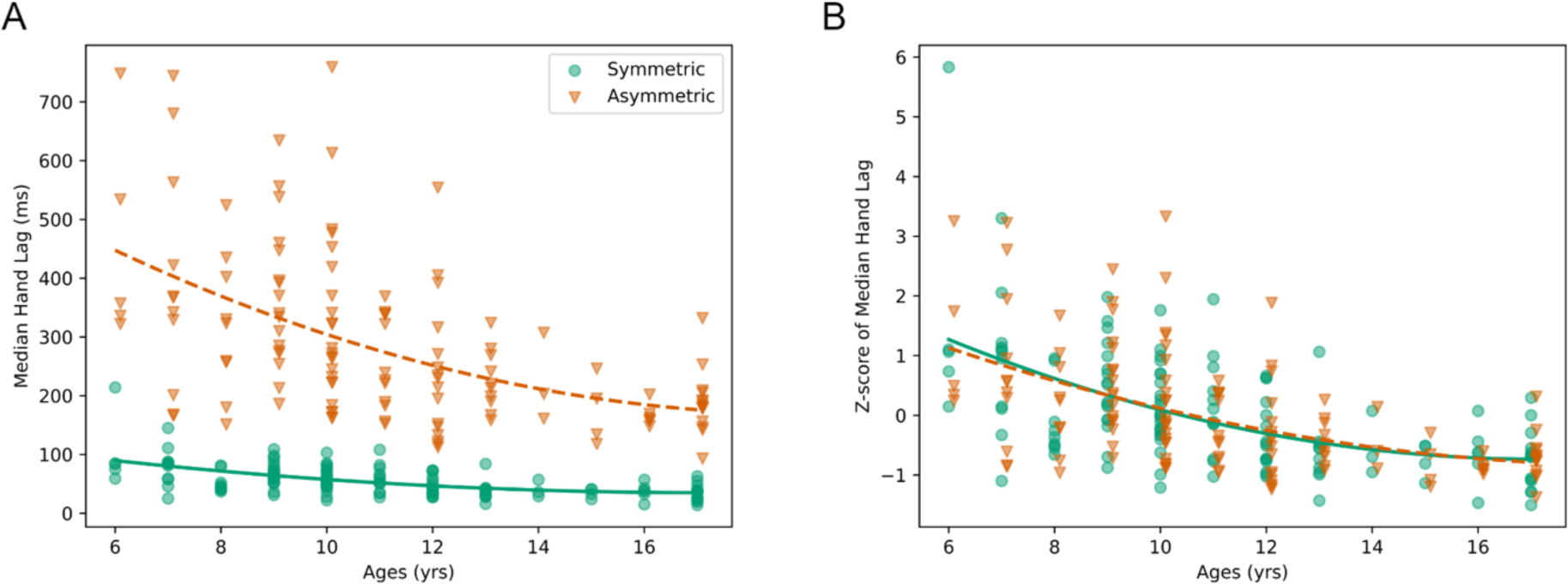
Development curves for hand lag. **A)** Median hand lag for the symmetric reaching task (green circle) and the asymmetric reaching task (orange triangle) for each participant is plotted against ages, with quadratic curves fit for each task (symmetric task, solid line; asymmetric task, dashed line). **B**) Z-score normalized median hand lags for each task are shown in order to compare tasks with different scales of hand lag. Darker colors indicate overlapping data points in both.

Comparing performance between the two tasks, we found that participants performed significantly better in the symmetric task than in the asymmetric task (*t(132)* = −28.713, *p* < 0.001). The difference in task performance between the symmetric reaching task and the asymmetric reaching task also stays consistent across ages (*R^2^* = 0.007, *p* = 0.347), which can also be observed by the parallel curves in Figure 3A.

Hand lag was significantly lower in the symmetric task compared to asymmetric reaching task (*t(129)* = 0.0, *p* < 0.001). The difference in median hand lag between the two tasks decreases across the ages (*R^2^* = 0.20, *p* < 0.001), with older adolescents having more similar median hand lag between the two tasks than younger children. However, the development of increasing synchrony between the hands (decreasing hand lag) is occurring in parallel, as shown by the normalized z-scores of hand lag in Figure 4B. Additionally, we found that participants reached the target with the dominant hand first in an average of 41.3% of the time for the symmetric task and 50.3% of the time in the asymmetric task.

### 3.3 Gender Effects on Performance

We found that without controlling for age, female participants outperformed male participants in the asymmetric task (*t(127) =* −2.398, p = 0.018) but not the symmetric task (*t(127) =* −1.594, p = 0.113). Further analysis using a two-way ANOVA shows that gender and age both have a significant effect on task performance in both tasks (symmetric reaching task gender: *F(1, 119) =* 4.719, *p* = 0.032, age: *F(4, 119) =* 28.999, *p* < 0.001; asymmetric reaching task gender: *F(1, 119) =* 11.304, *p* = 0.001, age: *F(4, 119) =* 31.793, *p* < 0.001) but there was not a significant interaction effect (symmetric reaching task *F(4, 119)* = 0.605, *p* = 0.660; asymmetric reaching task *F(4, 119)* = 0.878, *p* = 0.479). Examining gender differences in each age group, we found statistically significant differences, with females having higher scores than males, within the 10-11 year old age group in both tasks (symmetric task: *t(32) =* −2.107, *p* = 0.043, Cohen’s *D* = 0.707; asymmetric task: *t(32) =* 2.429, *p* = 0.021, Cohen’s *D* = 0.791) and within the 14-17 year old age group for the asymmetric task (*t(23) =* 2.223, *p* = 0.036, Cohen’s *D* = 1.051). As gender-based analyses were exploratory, multiple comparison corrections were not performed. Figure 5. illustrates the differences in performance between male and female participants in each age group.

**Fig. 5.**
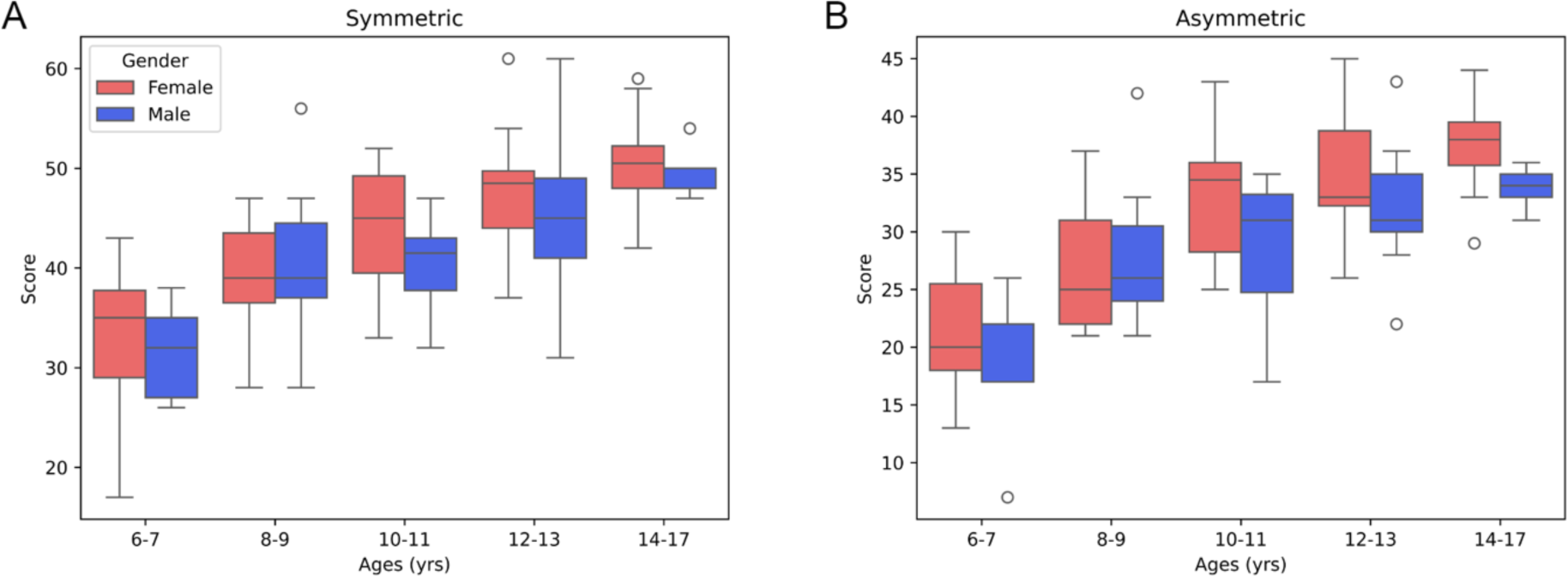
Gender differences across age groups. For participants within each age group, a box plot is drawn each for the self-reported female (left, red) and male (right, blue) participants. (**A**) Box plots for the symmetric reaching task are illustrated; gender differences were found for the 10-11 year olds. (**B**) Box plots for the asymmetric reaching task are illustrated; gender differences were found for the 10-11 year olds and 14-17 year olds.

### 3.4 Effect of Video Game Experience on Performance

One-way ANCOVAs with age as a covariate revealed that there was not a statistically significant difference in task performance between game play frequency groups (symmetric reaching task *F(5,132)* = 0.547, *p* = 0.701; asymmetric reaching task *F(5,132)* = 0.967, *p* = 0.428) or motion capture experience levels (symmetric reaching task *F(5,132)* = 0.744, *p* = 0.528; asymmetric reaching task *F(5,132)* = 1.736, *p* = 0.163). These effects are illustrated in Figure 6.

**Fig. 6.**
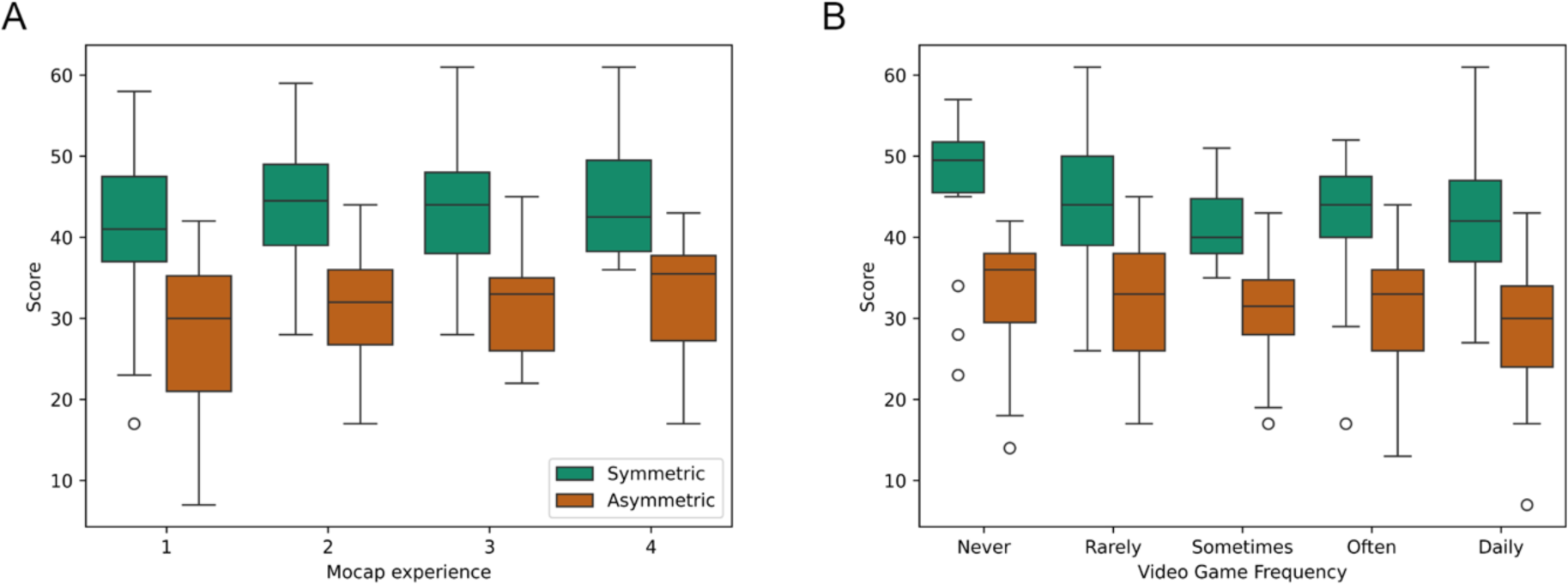
Performance Across Video Game Experience. Self-reported frequency the participant plays video games (**A**) and participant experience with motion capture (**B**) (1 being least experienced, 5 being most experienced) is plotted against the participant’s score in the symmetric reaching task (left, green) and the asymmetric reaching task (right, orange).

## 4. Discussion

This study used a large cohort of typically developing children to determine the development of bilateral reaching abilities using an assessment that combined computer vision with augmented reality gaming. Overall, we found that performance improved on both the symmetric and asymmetric reaching tasks with age, and children consistently performed better on the symmetric task than asymmetric. Development on both tasks occurred in parallel. We found that development did not plateau but rather continued to improve at a slower rate throughout adolescence. Lastly, we found that task performance was not related to prior video game experience, indicating that our tasks assess motor ability rather than gaming experience.

Performance on both the symmetric and asymmetric tasks improved with age, with age accounting for over 50% of the variance in score. While the greatest amount of change occurred before age 12, we found that improvements continued past age 12 and may not plateau by 17 years. This supports previous findings of continued development through adolescence (Boer et al., 2012), which matches the developmental course of the corpus callosum (Koerte et al., 2009, 2010). Interestingly, despite the asymmetrical task being more challenging (indicated by consistently lower scores than the symmetric task), development on the two tasks in terms of overall score occurred in parallel. We had expected the symmetric task to mature earlier based on the innate preference for symmetric movements. Our results indicate that this preference may be reflected in greater ease in performing symmetric reaches compared to asymmetric, however, both movements continue to mature throughout adolescence. While not reaching specific, the parallel development does match a prior finding that in-phase and anti-phase coordination developed at similar rates (Boer et al., 2012).

Hand lag was used as a measure of coordination reflecting the temporal synchrony in the reaches between the limbs. Similar to improvements in score, hand lag decreased with age, with age explaining nearly a third of the variability in hand lag. In addition to decreasing hand lag with age, variability across children also decreased. Children 12 years and younger varied widely, with some young children demonstrating hand lags on par with adolescents, and others significantly higher. By 13, the variability had decreased, and children were consistently performing with more synchronized movements on both symmetric and asymmetric tasks. The variability could be due to variability in brain development (Brown, 2017) or due to differences in experience, for instance, some children may engage in more tasks such as sports or crafts that require higher degrees of coordination and thus afford more practice (Dapp et al., 2021).

We found that females performed better than males on both the symmetric and asymmetric reaching tasks. When we further examined gender differences in each age group, we found the largest differences in the 10–11-year-olds for both tasks and the 14-17 year olds for the asymmetric tasks, with females outperforming males in both age groups. The developmental course of the corpus callosum can explain our age and gender-based findings.

In addition to continued growth and development of the corpus callosum across childhood and adolescence, puberty specifically has been associated with callosal growth (Chavarria et al., 2014). The apparent fiber density of the corpus callosum has been found to be related to age, sex, and puberty, with females having higher fiber density (Genc et al., 2018). The effect of puberty on corpus callosum development and fact that girls undergo puberty earlier than boys may explain why 10–11-year-old females outperformed males. Additional work has demonstrated that the maturation patterns in males and females and across different sections of the corpus callosum are different. Specifically, there have been two critical points at which the growth pattern of the corpus callosum shifts-first at 9-10 in girls and 11-12 in boys, and later at 15-16 in girls and 17-18 in boys (Luders et al., 2010). The earlier critical period for girls may explain their superior performance in our 10-11 year and 14-17 year age groups. We also note that our 14-17 year old group had more females than males and this finding would need to be replicated in a study specifically powered to examine gender differences at different age brackets. Additionally, the demographic information we collected was about gender, and while our findings match what is known about biological sex differences in development, we cannot determine if the differences we found between males and females were gender or sex-based differences.

This study had several limitations. First, our age and gender distributions were not equal. While conducting this study at the Minnesota State Fair allowed us to recruit a more representative sample than individuals who would typically participate in lab-based research, it reduced our control on the participants we recruited. The unequal distributions may have influenced our results on both age and gender-based differences. A larger sample across all ages and males and females would allow us to confirm our findings of gender differences. A longitudinal study with multiple time points would also further confirm our findings beyond our cross-sectional study. Additionally, while we attribute gender differences to differences in the onset of puberty, we did not assess Tanner stages to make definitive findings about the role puberty plays on bilateral coordination. We also used self-report to exclude individuals with motor impairments, however, were not specific in whether children with disabilities such as ADHD, autism, or developmental coordination disorder were excluded as it depended on whether the child/parent viewed them as having motor impairments. While we excluded individuals who were identified as outliers or who could not follow study instructions, it is likely our sample included children with subtle motor impairments that are prevalent in pediatric populations.

There were also limitations in terms our use of computer vision and game design. The use of marker-less capture with a single camera may have reduced tracking accuracy, especially in terms of motion in the direction of the camera. Individuals took different movement strategies with some limiting their reaching within the frontal plane and others reaching toward the screen (which was not captured). Since individuals had a practice period to become accustomed to playing the game, and we found no correlation with prior video game experience with performance, we do not believe variations in movement strategy (i.e. out of plane reaching) influenced overall performance. While it is challenging to quantify the overall accuracy of computer vision, our metrics and game were only reliant on the position of the hand in the frontal plane, which is less prone to error than joint angles, especially in multiple planes. Our game was also not designed to examine the full reaching kinematics during bilateral movements. This would require multiple camera angles to accurately capture joint angles. Characterizing kinematics would also call for greater standardization of target positions including angles and distances, and longer hold times at the targets.

Bilateral coordination is challenging to assess due to the variable nature of bilateral tasks. Our protocols measured bilateral reaching to two separate targets, which may not transfer to tasks where the arms reach to the same object such as a box. Future research would also be needed to determine how our gamified computer vision-based assessment would translate to performance on functional bilateral tasks. It can also be challenging to distinguish bilateral coordination from unilateral abilities, for instance, are the improvements with age just due to children getting faster at moving each arm rather than specific improvements in bilateral coordination. A unilateral task would also allow us to examine differences between unilateral and bilateral reaching development. Unfortunately, the unilateral task that we collected along with the two bilateral tasks was not designed as a direct comparison. Rather, we were assessing which hand participants chose to reach for a single target with. Since we did not control which hand participants reached with, we found high variability in whether they switched hands or used only one. A task in which participants can only use one hand would be needed to answer questions about how unilateral reaching ability relates to bilateral reaching.

The use of augmented reality games and computer vision to assess bilateral reaching has significant implications for clinical populations. The assessments used can provide a fast and inexpensive method to quantitatively assess bilateral reaching abilities. Collecting data in the environment of the Minnesota State Fair also demonstrates that the proposed methodology can easily be transferred to the clinic or bedside. It is also likely that this could be translated into a telehealth assessment. The quantitative nature can provide an objective measure that can be used to identify if a child has impairments relative to age-based norms, or track performance over time and in response to an intervention. The assessment (apart from demographic surveys) took less than five minutes to complete including setup and requires no specialized equipment besides a computer with a monitor and webcam. This could therefore provide both a means of assessing bilateral coordination which we currently lack, and quantitative markers of impairment which most clinical assessments of motor abilities do not provide. In the future, additional metrics using the kinematics can also be analyzed to further examine temporal and spatial coordination, however multiple cameras or a customized pose estimation algorithm may be necessary to reach the granular accuracy levels typically used in movement analysis. Lastly, the games could be adapted to be an intervention to address bilateral coordination, by providing high repetitions of practice in an enjoyable gamified manner.

Future research will further expand the sample size to be powered to address gender differences across our age continuum. We will also increase our upper age limit to determine at which point development plateaus on the bilateral tasks. We plan to expand the number of tasks to include unilateral reaching as well as cooperative reaching tasks. We will also analyze additional kinematics to determine specific movement strategies and how these may mature across development. Lastly, we will apply the protocol to children with motor impairments including cerebral palsy, developmental coordination disorder, and autism to determine if we can detect impairments in bilateral reaching abilities in these populations using objective measures. We may need to adapt the protocol for clinical populations to account for slower movement speeds and additional fatigue.

## Acknowledgements

This work was supported by the Graduate Training Program in Sensory Science: “Optimizing the Information Available for Mind and Brain”: Grant No.: DGE-1734815 and NIH P2C HD10191301.

## CRediT Author Statement

**Shelby Ziccardi:** conceptualization, methodology, software, formal analysis, investigation, writing-original draft, writing-review and editing, visualization. **Samantha Timanus**: methodology, software. **Ghazaleh Ashrafzadehkian**: methodology, software, investigation. **Stephen Guy:** conceptualization, methodology, investigation, writing-review and editing, supervision. **Rachel Hawe:** conceptualization, methodology, investigation, writing-original draft, writing-review and editing, supervision, funding acquisition.

